# Hepatocyte and adipocyte CDO1-mediated intracellular cysteine catabolism differentially modulates diet-induced obesity and fatty liver in mice

**DOI:** 10.64898/2025.12.05.692669

**Authors:** Jianglei Chen, Yung-Dai Clayton, Yanhong Du, David Matye, Lijie Gu, Mohammad Nazmul Hasan, Tiangang Li

**Author notes:** **Correspondence** Tiangang Li, PhD Department of Biochemistry and Physiology Harold Hamm Diabetes Center University of Oklahoma Health Sciences Center Oklahoma City, OK 73104.

## Abstract

**Background and Aims:** Plasma cysteine has been positively linked to BMI in humans. Consistently, dietary cysteine restriction significantly decreases obesity in mice via mechanisms that are incompletely defined. Cysteine dioxygenase type 1 (CDO1) mediates the major cysteine catabolism pathway and is highly expressed in hepatocytes and adipocytes. The goal of this study is to determine the impact of this endogenous cysteine catabolism pathway on obesity and fatty liver disease.

**Methods:** Diet-induced obesity and fatty liver disease were studied in hepatocyte-specific CDO1 knockout mice (L-CDO1-KO), hepatocyte-specific CDO1 transgenic mice (L-CDO1-Tg), adipocyte-specific CDO1 knockout mice (Ad-CDO1-KO), and adipocyte-specific CDO1 transgenic mice (Ad-CDO1-Tg).

**Results:** Deletion of hepatocyte CDO1 decreased cysteine conversion to cysteine sulfinic acid, and had modest impact on liver cysteine, GSH, or taurine abundance. When fed a Western diet (WD), L-CDO1-KO mice showed elevated liver injury markers and inflammatory infiltration independent of obesity or steatosis. When fed a fibrogenic high fat/cholesterol/fructose diet (HFCFr), L-CDO1-KO mice developed worsened liver fibrosis. In contrast, L-CDO1-Tg mice fed WD showed lower blood glucose, but similar degree of obesity and steatosis compared to WT mice. Deletion of CDO1 in white and brown adipocytes of Ad-CDO1-KO mice had no effect on WD-induced obesity. In contrast, overexpression of CDO1 in white and brown adipocytes attenuated WD-induced obesity, which resulted in reduced hepatic steatosis.

**Conclusion:** Genetic manipulation of intracellular cysteine catabolism in hepatocytes and adipocytes differentially modulates diet-induced obesity and fatty liver, which warrants future mechanistic investigation to better understand cellular cysteine sensing mechanisms and cysteine control of cell metabolism.

## Introduction

Cysteine is a non-essential sulfur amino acid that can be synthesized from methionine via the transsulfuration pathway (1). Liver is the major sulfur amino acid metabolism organ and expresses high level of methionine and cysteine metabolizing enzymes. Cysteine is the substrate for synthesis of protein, glutathione (GSH), taurine, coenzyme A, pyruvate, and H_2_S (1), which support various cellular functions including anti-oxidant defense, bile acid conjugation, fatty acid oxidation, and energy metabolism (**Fig 1A**) (2, 3). Cysteine is irreversibly metabolized by the cysteine dioxygenase type 1 (CDO1) to cysteine sulfinic acid, which is further converted to taurine for bile acid conjugation (4).

**Figure 1.**
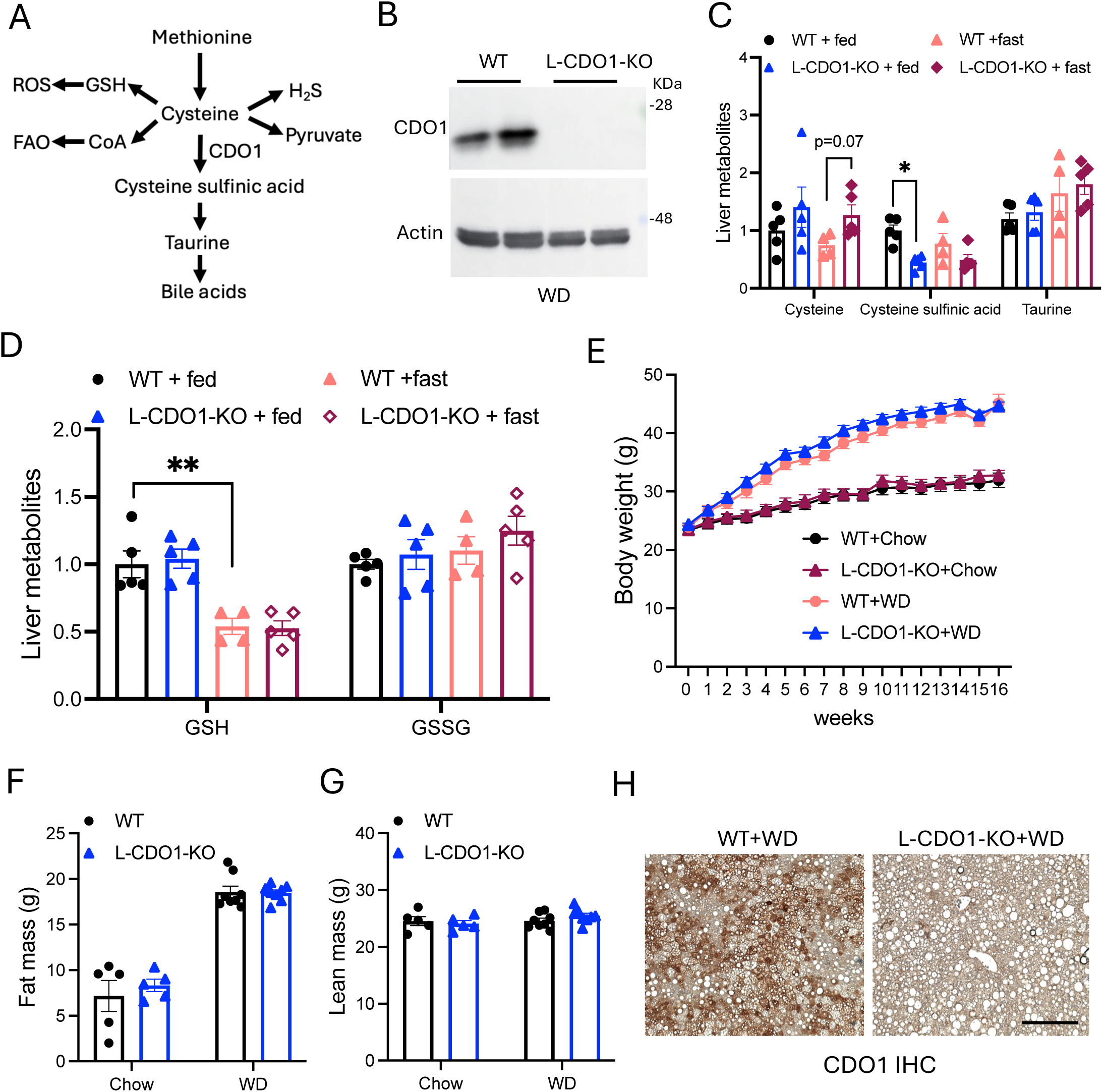
Hepatic CDO1 knockout does not affect 16 weeks Western diet-induced obesity. **A.** Cysteine metabolism pathways. CoA: coenzyme A. FAO: fatty acid oxidation. **B.** Liver CDO1 protein. **C, D.** Liver metabolites. n=4-5. Mice were fasted overnight (fast) or fed chow for 6 hours after overnight fasting (fed). **E-H.** Mice were fed chow or WD for 16 weeks. n=5-8. Body weight (E), body composition (F, G), and representative of immunohistochemistry stain of CDO1 in liver section. Scale bar = 250 μm. All results are shown as mean ± SEM.

Hepatocyte CDO1 knockout elevated intracellular cysteine concentration (5). CDO1 protein is feedforward stabilized by cysteine (6). In addition, excessive cysteine is incorporated into the GSH pool. These mechanisms prevent intracellular cysteine accumulation that could lead to a buildup of cysteine metabolites and cytotoxicity in various organs (7, 8). We have previously reported that bile acids inhibit hepatic CDO1 to coordinate the de novo synthesis of bile acid and taurine in hepatocytes (9). Hepatic CDO1 induction as a result of attenuated bile acid repression decreases hepatic cysteine availability, which can impair hepatic GSH regeneration capacity under oxidative stress and exacerbate acetaminophen hepatotoxicity (9). In contrast, hepatic CDO1 knockout preserves cysteine to promote GSH synthesis and protects against acetaminophen hepatotoxicity (10). CDO1 is often silenced in cancer cells, which provides a survival advantage by preserving cellular cysteine pool and GSH synthesis (11). Furthermore, hepatic cysteine is a key substrate in the synthesis of coenzyme A, which needs to be upregulated to support fatty acid oxidation under conditions of increased hepatic fatty acid influx (3, 12). Fatty liver disease is associated with chronic oxidative stress among other pathogenic drivers. Whether reduced hepatic CDO1 function is associated with beneficial outcomes in fatty liver disease is still not known.

It has been recognized years ago that circulating cysteine concentration positively correlates with body mass index (BMI) and obesity in humans (13–15). Dietary sulfur amino acid restriction causes weight loss in both humans and animal models (16). Cysteine supplementation fully reversed the weight loss effect of dietary methionine restriction, which provided experimental evidence that the weight loss effect of dietary methionine restriction is largely attributed to cysteine deficiency (17, 18). Consistently, we have recently reported that dietary protein restriction only decreased hepatic cysteine abundance without causing an overall amino acid deficiency (3), and dietary cystine supplementation fully abolished the weight loss effect of dietary protein restriction (19). Recent studies have attributed the weight loss effect of dietary cysteine deficiency to both metabolic reprogramming in liver and adipose tissues and altered sympathetic neuronal activity (20, 21). However, the complex mechanisms mediating the robust weight loss effect of dietary cysteine restriction are still incompletely understood. Global CDO1 knockout mice showed elevated circulating cysteine and altered energy metabolism (22), which suggests that CDO1-mediated cysteine catabolism may not only regulate intracellular cysteine abundance but possibly systemic cysteine exposure. Given the significant impact of dietary cysteine intake on obesity development, whether changes of endogenous cysteine catabolism modulate diet-induced obesity warrants investigation.

Global CDO1 knockout in mice was associated with multi-organ toxicity and early mortality (7, 8), which prevents further evaluation of the metabolic impact of the CDO1-mediated cysteine catabolism in organ specific manners. In this study, we generated mouse models with CDO1 knockout or overexpression selectively in hepatocytes or adipocytes, two organs that express relatively high levels of CDO1 and are important in energy metabolism and obesity development. Our results revealed that genetic manipulation of hepatic and adipocyte cysteine catabolism via CDO1 differentially affected the susceptibility to diet-induced obesity and fatty liver disease progression.

## Results

### Hepatocyte CDO1 knockout mice (L-CDO1-KO) fed a Western diet (WD) showed exacerbated liver inflammatory infiltration and injury

AAV-TBG-cre injection effectively deleted CDO1 in CDO1-flox mice (**Fig 1B**). It was noted that hepatic cysteine sulfinic acid was decreased by ∼50% instead of being absent upon CDO1 knockout (**Fig 1C**). CDO1 is the only enzyme know so far to mediate cysteine catabolism to cysteine sulfinic acid. The source of liver cysteine sulfinic acid in L-CDO1-KO mice is unclear and could be non-enzymatic oxidation of cysteine or of none-hepatocyte origin since whole liver lysates were used for such measurements. Under fed condition, liver cysteine was similar between WT and L-CDO1-KO mice. Under fasting condition when liver cysteine was lower, knockout of hepatocyte CDO1 resulted in elevated liver cysteine as we reported previously (10). However, without oxidative stress, hepatic CDO1 knockout did not affect basal liver GSH, GSSG, or taurine in either fasting or fed condition in chow-fed mice (**Fig 1C**).

When fed a chow or WD for 16 weeks, L-CDO1-KO mice show similar weight gain and adiposity as WT mice (**Fig 1E-G**). Immunohistochemistry stain of CDO1 confirmed that liver CDO1 was absent in L-CDO1-KO mice after 16 weeks of WD feeding (**Fig 1H**). Upon evaluation of liver injury, we found that L-CDO1-KO mice fed WD showed elevated blood liver enzymes (**Fig 2A-B**), which was not due to increased hepatic steatosis (**Fig 2C-E**). However, L-CDO1-KO mice showed accumulation of larger lipid droplets (**Fig 2E**). There was also significantly higher number of F4/80 positive lipid droplet-associated macrophages that formed crown-like structures in L-CDO1-KO mice (**Fig 2F-G**). Consistently, L-CDO1-KO mice showed higher expression of MCP-1 and TNFα mRNA than WT mice when fed WD (**Fig 2H-I**). These results show that L-CDO1-KO mice were more susceptible to developing liver inflammatory infiltration and injury when fed WD.

**Figure 2.**
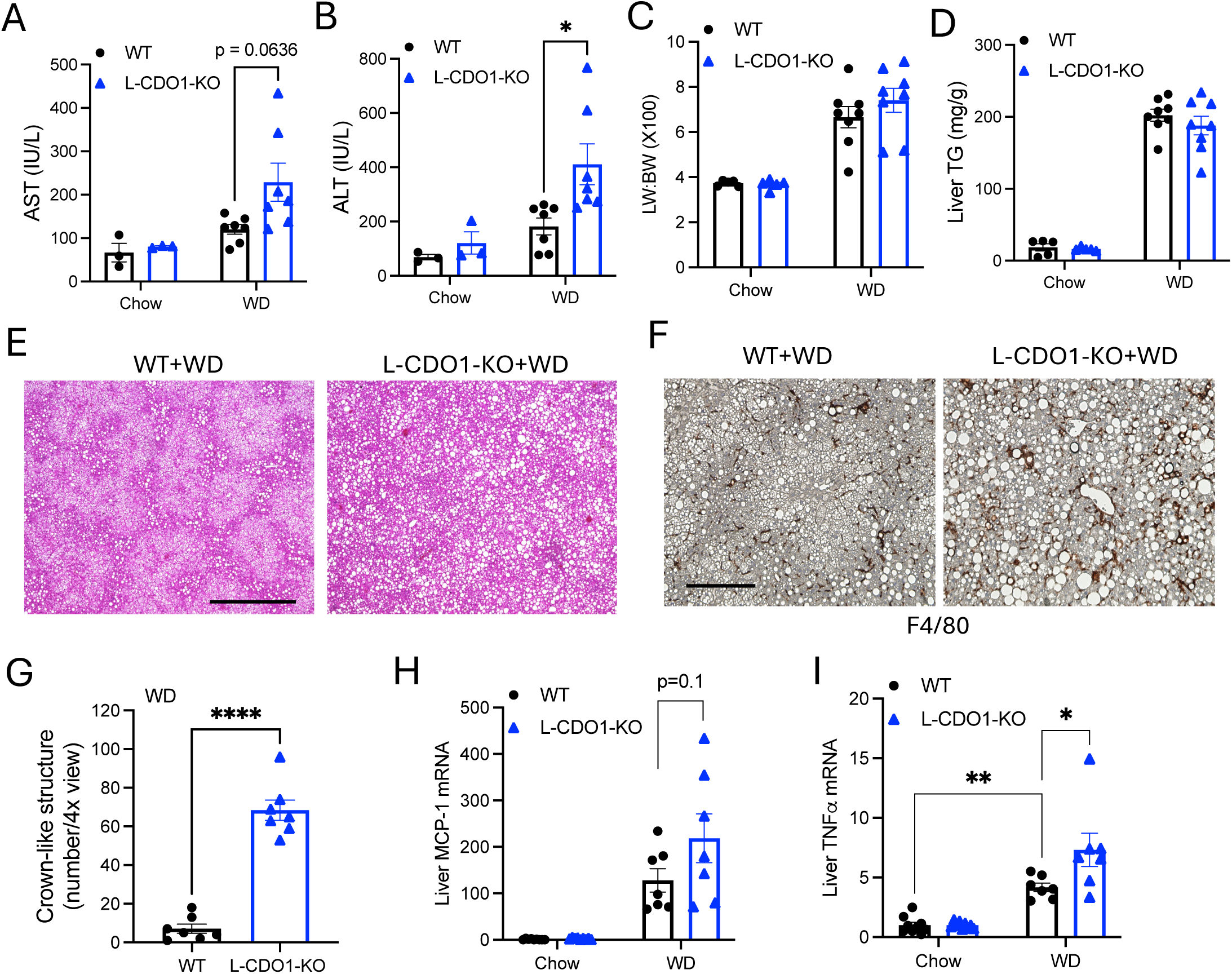
Hepatic CDO1 knockout mice show worsened liver injury and inflammation when fed Western diet for 16 weeks. Mice were fed chow or WD for 16 weeks. n=5-8. **A-B.** Blood aspartate aminotransferase (AST) and alanine aminotransferase (ALT). **C.** Liver weight (LW) to body weight (BW) ratio. **D.** Liver triglycerides (TG). **E.** Liver H&E stain. Scale bar = 600 μm. **F.** Immunohistochemistry stain of F4/80. Scale bar = 250 μm. **G.** Quantification of the number of F4/80 positive crown structure of “F”. **H-I.** Liver mRNA expression. MCP-1: Monocyte chemoattractant protein-1; TNFα: tumor necrosis factor α. All results are shown as mean ± SEM.

### L-CDO1-KO mice fed a fibrogenic high fat/cholesterol/fructose (HFCFr) diet developed worsened liver fibrosis

To further investigate how liver CDO1 deficiency affects the development of liver fibrosis, we fed WT and L-CDO1-KO mice a HFCFr diet, which we previously showed caused liver fibrosis in WT after 8 months of feeding (23). L-CDO1-KO mice fed HFCFr diet showed significantly higher weight gain and adiposity than WT mice (**Fig 3A-C**). Fasting blood glucose was also higher in L-CDO1-KO mice than WT (**Fig 3D**). L-CDO1-KO mice also showed modestly higher liver weight to body weight ratio and blood liver enzymes than WT mice. Liver CDO1 protein remained low in L-CDO1-KO mice after HFCFr diet feeding (**Fig 3H**). However, liver triglyceride and cholesterol levels were similar in L-CDO1-KO mice and WT mice (**Fig 3I-K**), but L-CDO1-KO mice developed more advanced fibrosis than WT mice (**Fig 3L**). We also analyzed a cohort of WT and L-CDO1-KO mice after 4 months of

**Figure 3.**
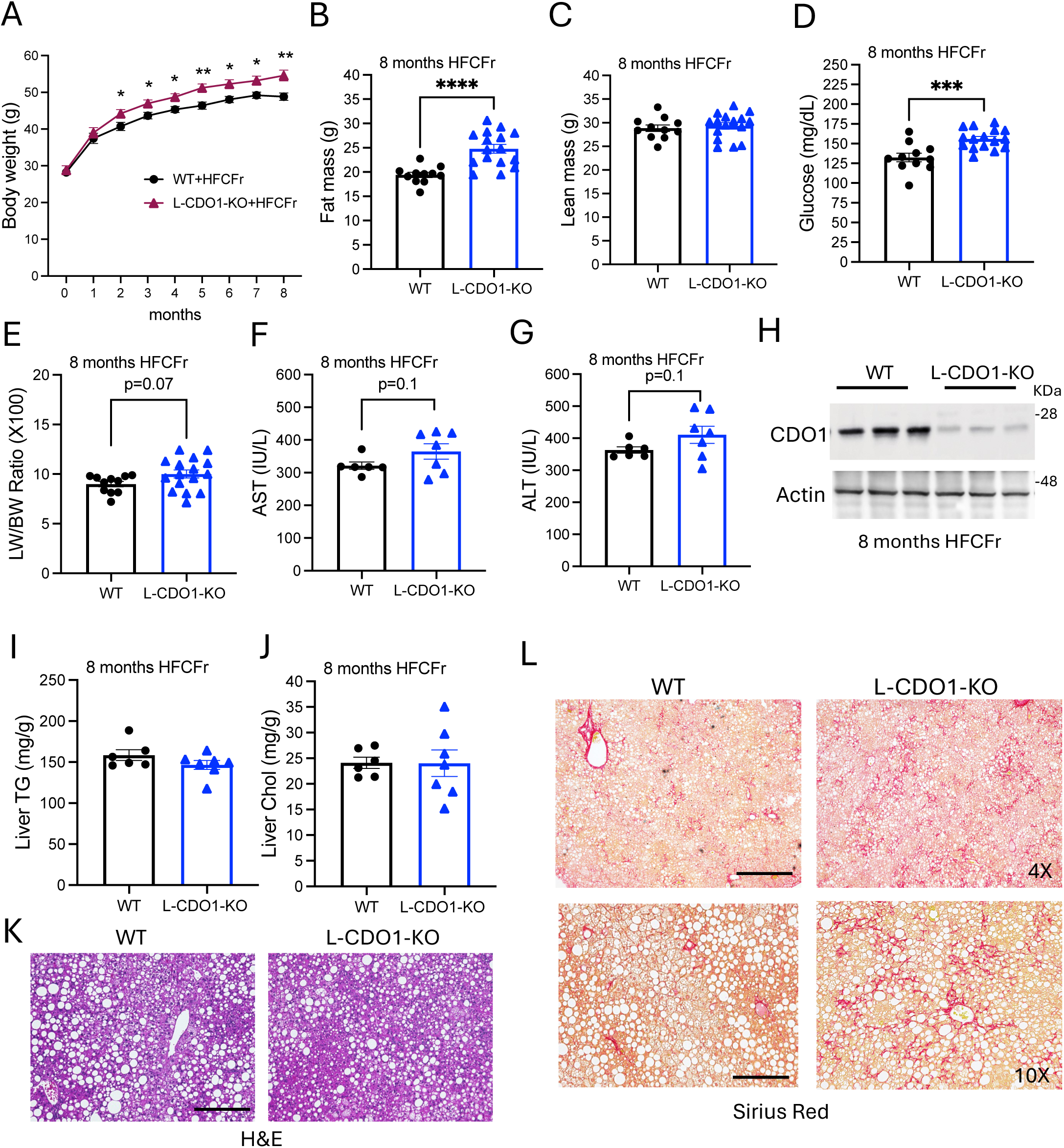
Hepatic CDO1 knockout mice show worsened liver fibrosis when fed high fat cholesterol fructose (HFCFr) diet for 8 months. Mice were fed HFCFr diet for 8 months. **A.** Body weight. n=11-15. **B-C.** Body composition. **D.** Blood glucose after 6 hours fasting measured during the 8^th^ month of HFCFr diet feeding. **E.** Liver weight (LW) to body weight (BW) ratio. **F-G.** Blood aspartate aminotransferase (AST) and alanine aminotransferase (ALT). **H.** Liver CDO1 protein after 8 months HFCFr diet feeding. **I.** Liver triglycerides (TG). **J.** Liver total cholesterol. **K.** Liver H&E stain. Scale bar = 250 μm. **L.** Liver Sirius red stain. Upper panels, 4x view, Scale bar = 600 μm. Lower panels, 10 x view, Scale bar = 250 μm. All results are shown as mean ± SEM.

HFCFr diet feeding. While L-CDO1-KO mice and WT mice showed similar hepatic triglyceride accumulation and both groups were absent of liver fibrosis after 4 months of HFCFr diet feeding (**Fig 4A-B**), L-CDO1-KO mice showed significantly higher blood liver enzymes and liver cytokine mRNA expression (**Fig 4C-E**), suggesting that worsened liver injury and inflammation is a likely cause of developing more advanced liver fibrosis at the later stage.

**Figure 4.**
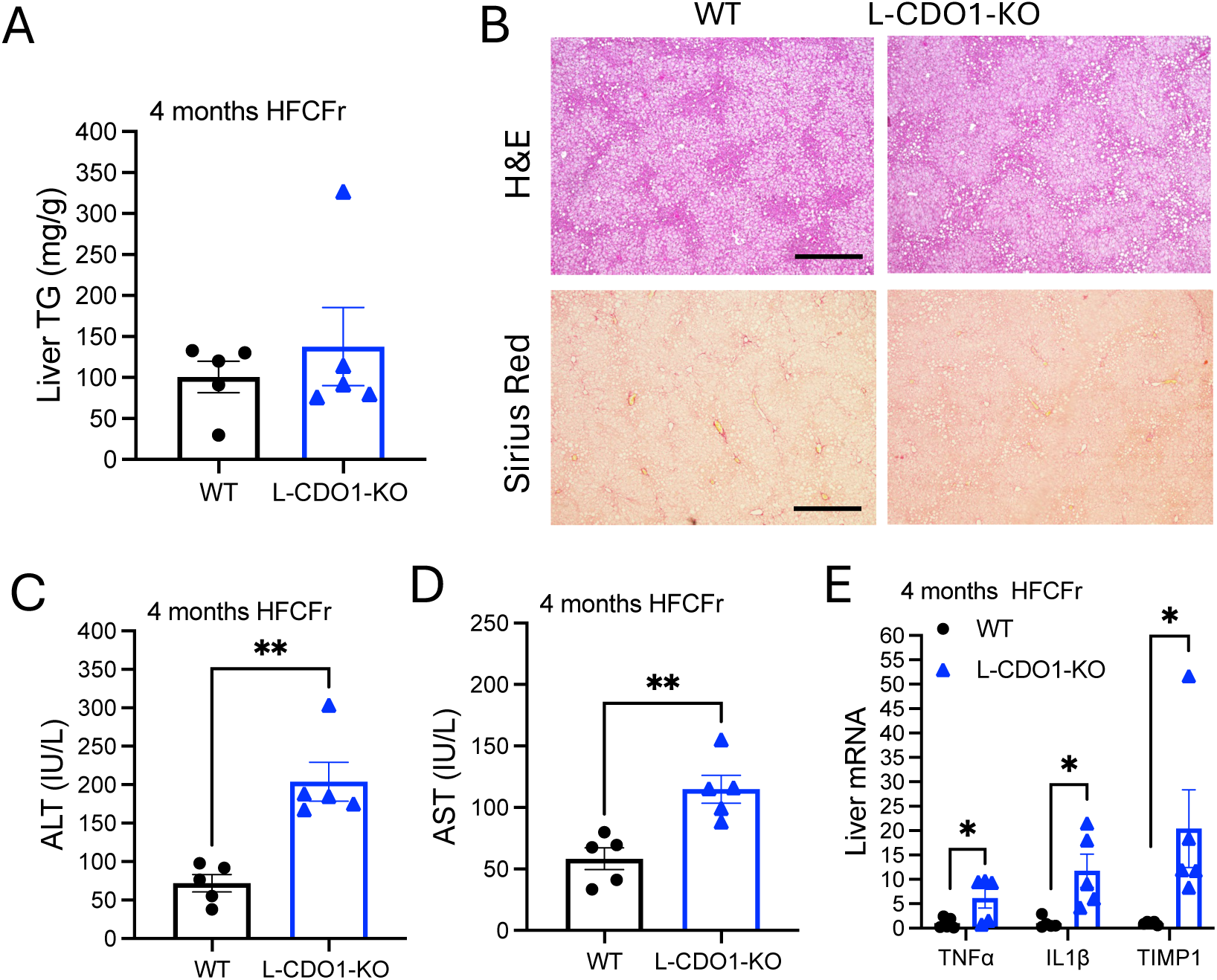
Hepatic CDO1 knockout mice show worsened liver injury and inflammation when fed high fat cholesterol fructose (HFCFr) diet for 4 months. Mice were fed HFCFr diet for 4 months. n=5. **A.** Liver triglycerides (TG). **B.** Liver H&E stain and Sirius red stain. Scale bar = 600 μm. **C-D.** Blood aspartate aminotransferase (AST) and alanine aminotransferase (ALT). **E.** Liver mRNA expression. TNFα: tumor necrosis factor α; IL-1β: interleukin-1β; TIMP-1: tissue inhibitor of metalloproteinase-1. All results are shown as mean ± SEM.

After 8 months of HFCFr diet feeding. L-CDO1-KO mice did not show altered cysteine but slightly decreased methionine, homocysteine and cystathionine levels (**Fig 5A**). L-CDO1-KO mice showed modestly but significantly increased liver GSH than WT mice (**Fig 5B**), which was consistent with increased hepatic GSH synthesis capacity under oxidative stress condition as we previously demonstrated in L-CDO1-KO mice challenged with acetaminophen overdose (10). L-CDO1-KO mice showed similar liver acylcarnitine and free fatty acids as WT mice (**Fig 5C-D**). L-CDO1-KO mice also showed similar level of liver taurine and taurine-conjugated bile acids (**Fig 5E-F**). Taken together, these results suggest that blocking hepatic cysteine catabolism by CDO1 knockout increased the susceptibility to hepatic injury and inflammation, which leads to the development of worsened liver fibrosis in mice fed WD or HFCFr diet.

**Figure 5.**
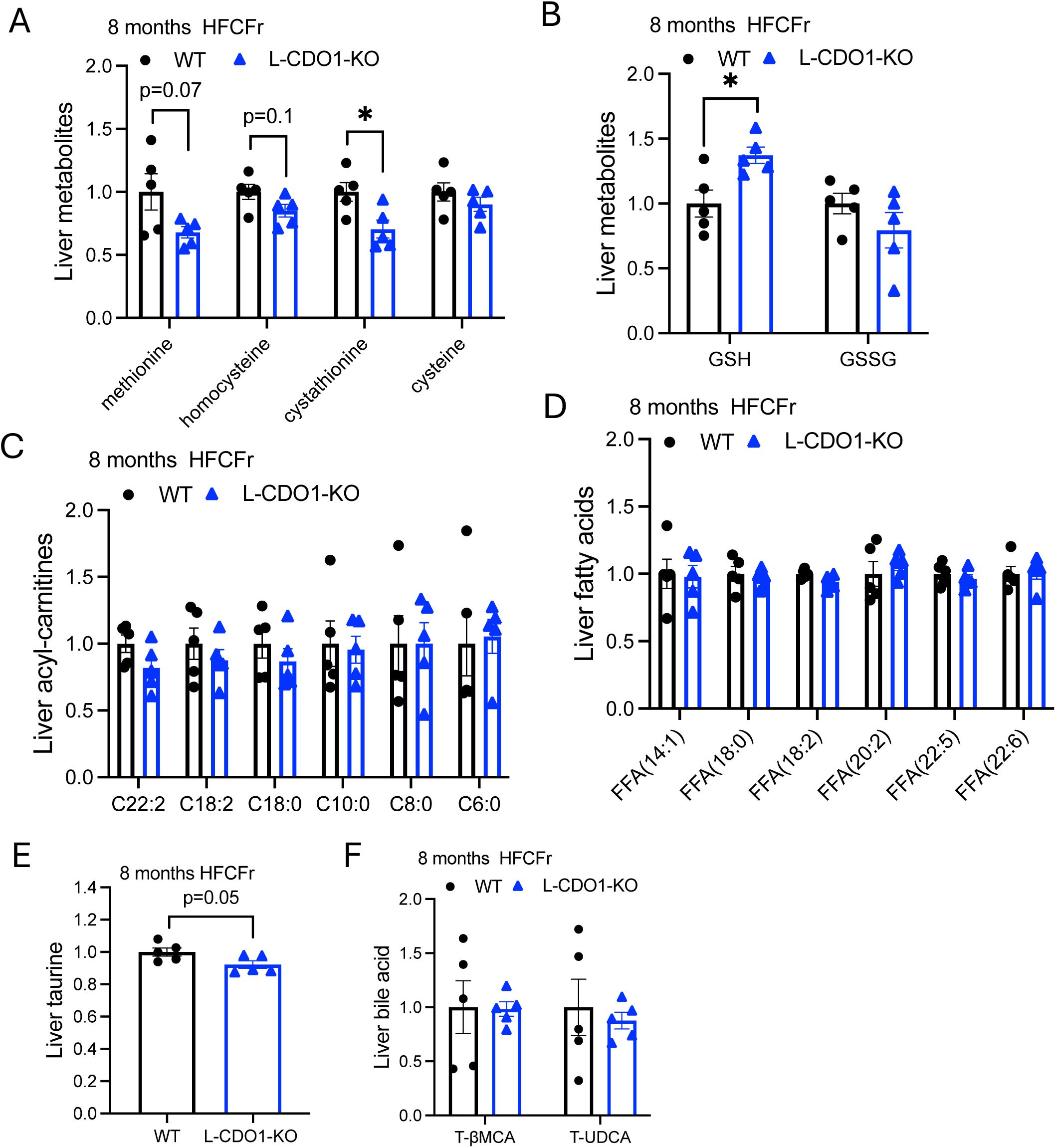
Liver metabolites in hepatic CDO1 knockout mice fed high fat cholesterol fructose (HFCFr) diet for 8 months. Mice were fed HFCFr diet for 8 months. n=5. Liver metabolites were measured by mass spectrometry. All results are shown as mean ± SEM. The average value of WT was set as “1”. T-βMCA: taurine-conjugated β muricholic acid; T-UDCA: taurine-conjugated ursodeoxycholic acid.

### Hepatic CDO1 overexpression lowered fasting glucose without affecting WD-induced obesity or hepatic steatosis

To further investigate the metabolic effects of hepatic CDO1 induction, we next generated hepatocyte CDO1 transgenic mice (L-CDO1-Tg) where injection of AAV-TBG-cre to CDO1(ROSA26) mice selectively induces hepatocyte CDO1 overexpression (**Fig 6A**). In a cohort of 2 weeks WD fed mice, we found that hepatic cysteine was modestly decreased while cysteine sulfinic acid was elevated, which reflected increased hepatic CDO1 activity (**Fig 6B**). However, hepatic CDO1 overexpression did not increase hepatic taurine or reduce hepatic GSH in mice (**Fig 6B-C**). When fed a WD, L-CDO1-Tg mice showed similar body weight and adiposity as WT mice (**Fig 6D-E**). Hepatic steatosis was also similar between L-CDO1-Tg mice and WT mice (**Fig 6F-G**). Fasting blood glucose was significantly lower in L-CDO-Tg mice than WT (**Fig 6H**). This finding was consistent with our previous report that bile acid sequestrant-mediated glucose lowering may be significantly attributed to hepatic CDO1 induction due to attenuated bile acid signaling in the liver (19). However, promoting hepatic cysteine catabolism decreases hepatic cysteine but does not attenuate obesity development.

**Figure 6.**
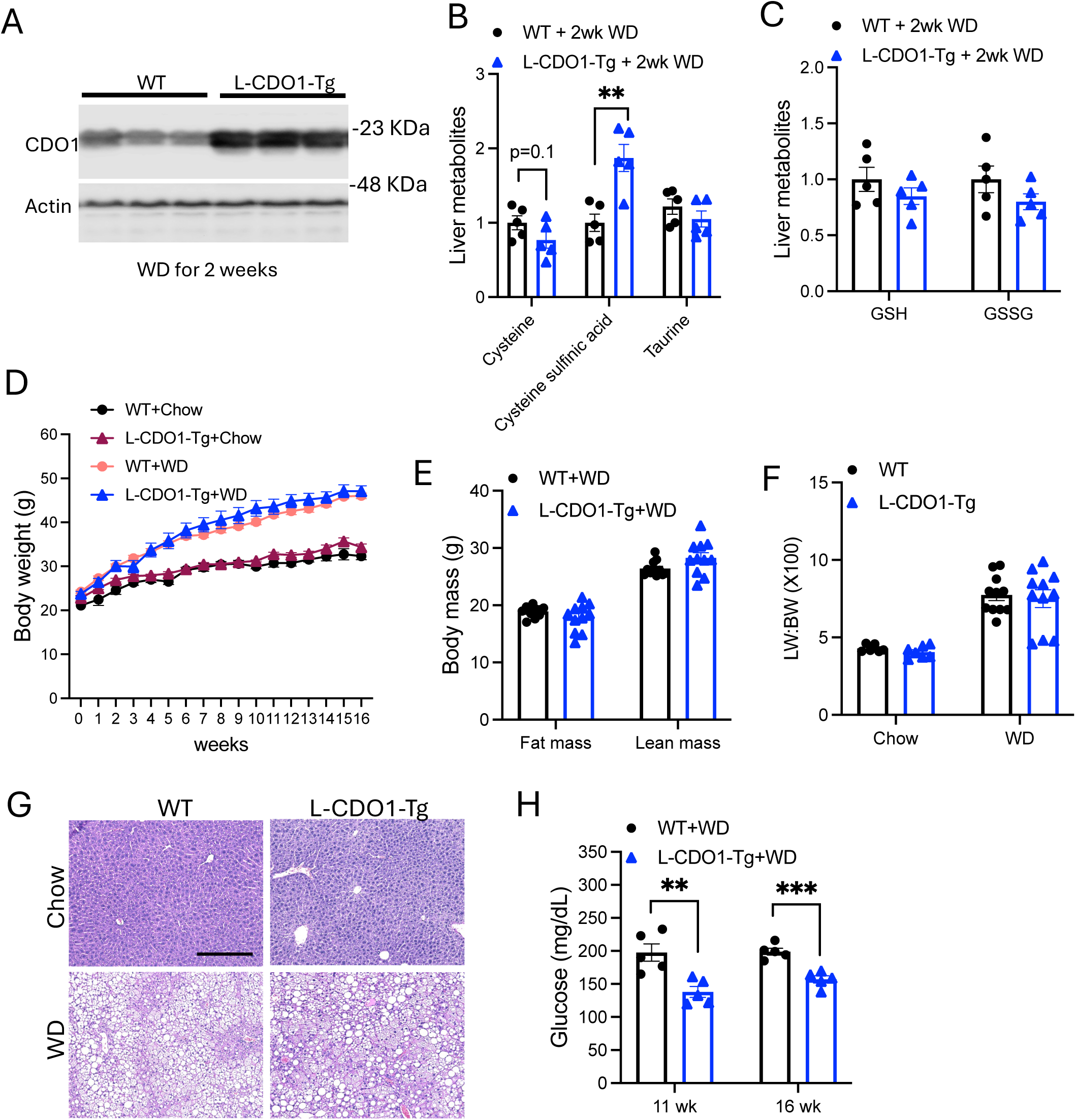
Hepatic CDO1 overexpression lowers blood glucose but does not affect 16 weeks Western diet-induced obesity. **A.** Liver CDO1 protein in mice fed WD for 2 weeks. **B-C.** Liver metabolites in mice fed WD for 2 weeks. n=5. **D-H.** Mice were fed chow or WD for 16 weeks. n=6-11. Body weight (D), body composition (E), LW:BW ratio (F), liver H&E stain, Scale bar = 250 μm (G), and 6 hours fasting blood glucose. All results are shown as mean ± SEM.

### Adipose CDO1 overexpression attenuates obesity and hepatic steatosis

We have recently shown that reducing cysteine concentration in culture medium dose-dependently inhibited adipogenesis in vitro (19), suggesting that dietary cysteine intake may act directly on adipocytes to modulate obesity development. In addition to hepatocytes, adipose tissue also shows high CDO1 expression. To determine if CDO1-mediated cysteine catabolism modulates obesity development, we developed adipocytes CDO1 knockout mice (Ad-CDO1-KO) that showed deletion of CDO1 in both white adipose and brown adipose (**Fig 7A**). Upon WD feeding, Ad-CDO1-KO mice show similar weight gain as WT mice (**Fig 7B**). White adipose and brown adipose of Ad-CDO1-KO and WT mice also showed similar histologic appearance (**Fig 7C-D**). Further metabolic characterization was not carried out in this cohort of mice.

**Figure 7.**
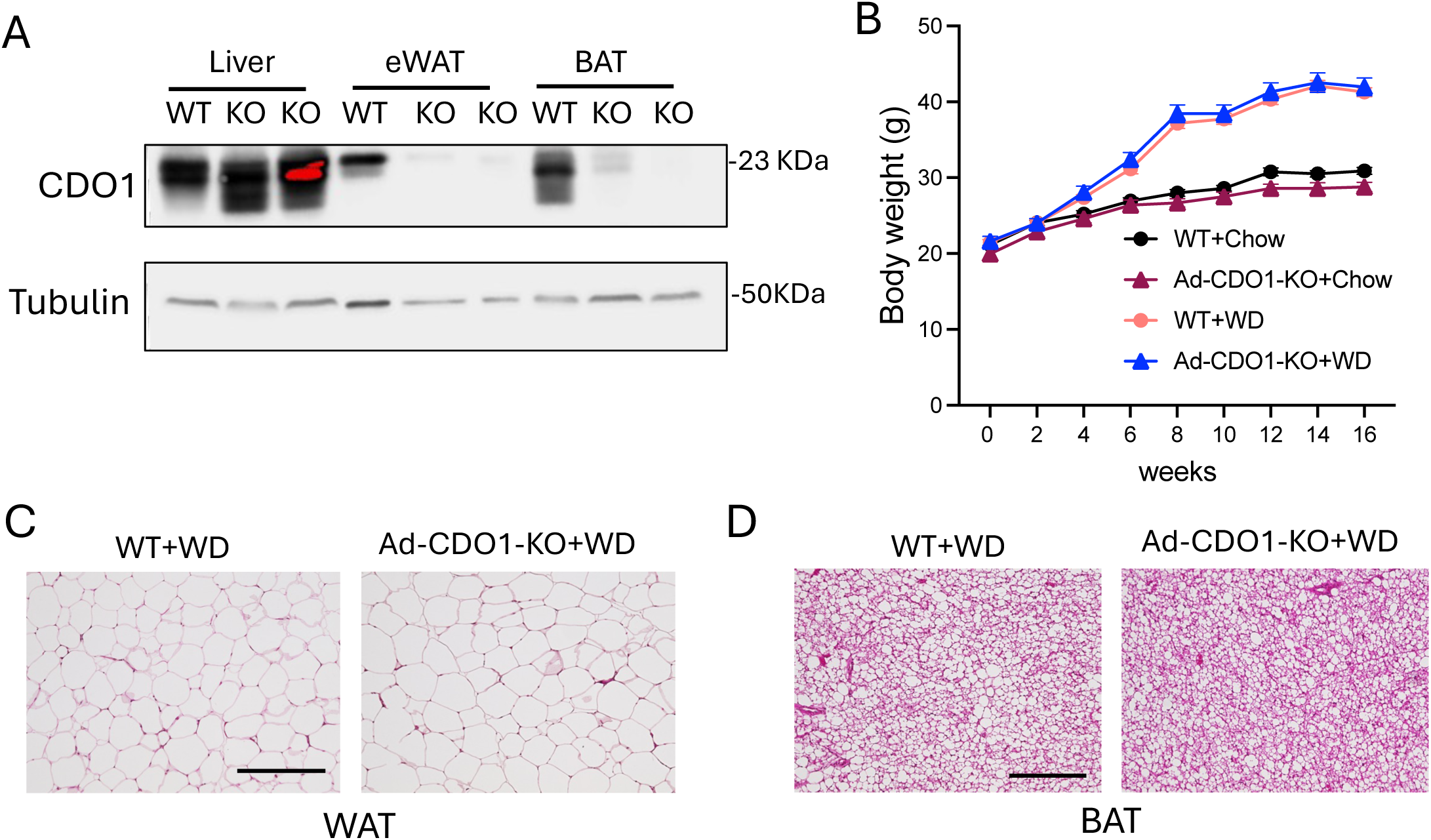
Adipocyte CDO1 knockout does not affect 16 weeks Western diet-induced obesity. **A.** CDO1 protein in liver, epidydimal white adipose tissue (eWAT), and brown adipose tissue (BAT) of chow-fed mice. **B-D.** Mice were fed WD for 16 weeks. n=7-17. Body weight (B), eWAT H&E stain, Scale bar = 250 μm (C), and BAT H&E stain, Scale bar = 600 μm (D). BW results are shown as mean ± SEM.

We next generated adipocyte CDO1 transgenic mice (Ad-CDO1-Tg) which showed increased CDO1 expression in both white adipose and brown adipose (**Fig 8A**). When fed a WD, Ad-CDO1-Tg mice showed significantly attenuated weight gain, which was attributed to decreased fat mass (**Fig 8B-D**). Consistently, white adipocytes of Ad-CDO1-Tg mice appeared to be smaller and brown adipose of Ad-CDO1-Tg mice showed less lipid droplet accumulation than WT mice (**Fig 8E**). Hepatic steatosis was also attenuated in WD-fed Ad-CDO1-Tg mice than WT mice (**Fig 8E-F**). Taken together, these results show that, while blocking adipocyte cysteine catabolism upon CDO1 knockout does not promote obesity development, activation of adipocyte cysteine catabolism via CDO1 induction attenuates adiposity and obesity in mice.

**Figure 8.**
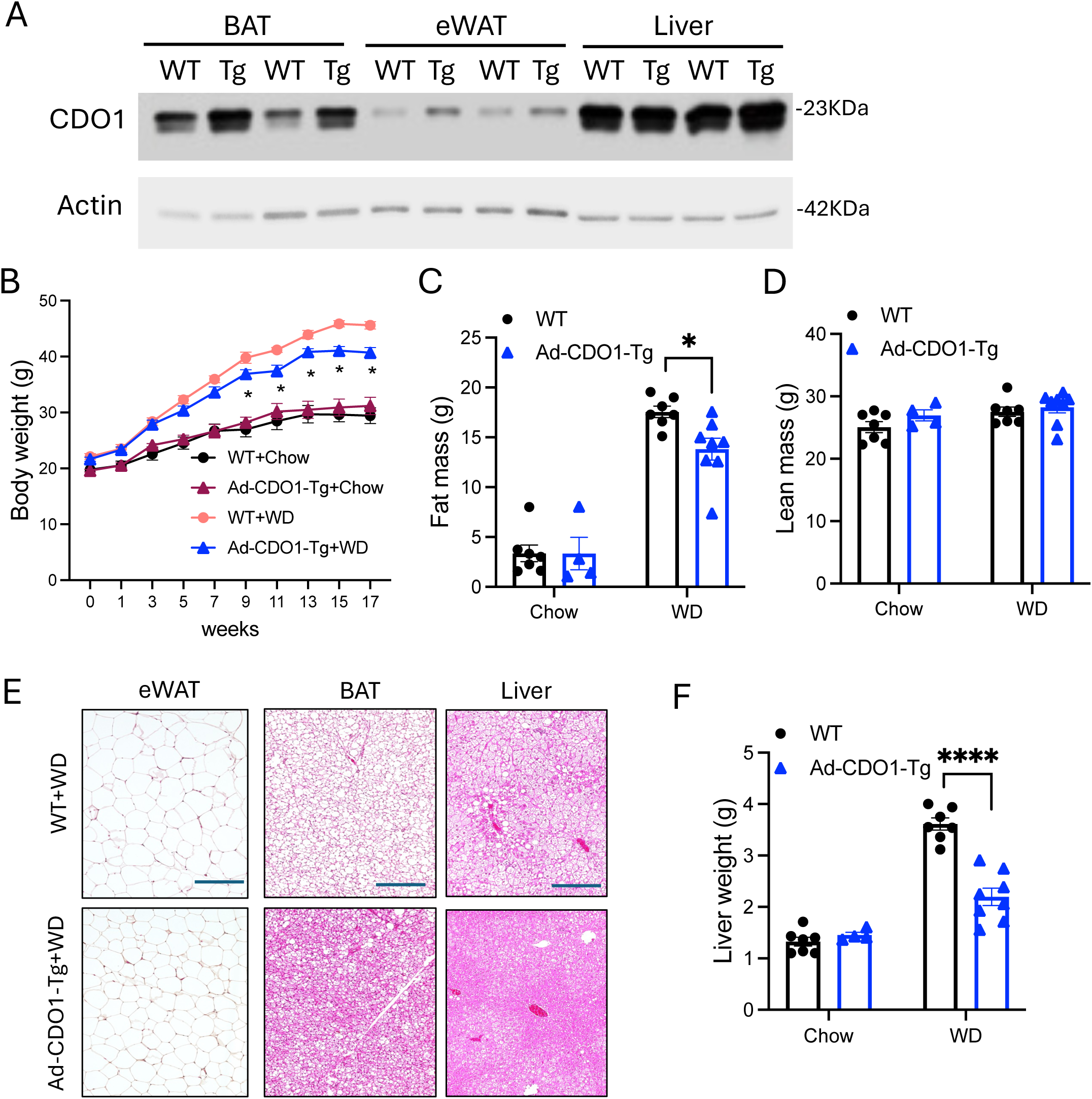
Adipocyte CDO1 overexpression attenuates Western diet-induced obesity and hepatic steatosis. **A.** CDO1 protein in liver, epidydimal white adipose tissue (eWAT), and brown adipose tissue (BAT) of chow-fed mice. **B-F.** Mice were fed chow or WD for 16 weeks. n=4-8. Body weight (B), body composition (C, D), and representative H&E stain of eWAT, BAT, and liver sections, Scale bar = 250 μm (E), and liver weight (F). All results are shown as mean ± SEM.

## Discussion

CDO1 critically controls cellular cysteine, which is utilized for hepatic synthesis of GSH and coenzyme A that support hepatic antioxidant defense and fatty acid oxidation, respectively. While blocking CDO1-mediated cysteine catabolism could preserve cysteine to enhance GSH regeneration during APAP hepatotoxicity where GSH depletion is a major driver of liver injury (10), deletion of liver CDO1 did not protect fatty liver where oxidative stress is only one of many pathogenic drivers of disease progression. In contrast, hepatic CDO1 KO promoted liver inflammatory infiltration and injury when mice were fed a WD. Consistently, this initial susceptibility to liver injury led to more advanced liver fibrosis when mice were fed the HFCFr diet for 8 months. Worsened liver inflammation and fibrosis were not due to higher accumulation of liver fat or lipotoxic lipid species including acyl-carnitine or fatty acids. Furthermore, deletion of CDO1 did not cause hepatic taurine deficiency or bile acid conjugation defect, excluding dysregulation of bile acid as an underlying cause. Because CDO1 is the only known enzyme that metabolizes cysteine to cysteine sulfinic acid, deletion of CDO1 likely caused a marked blockade of cysteine elimination in the liver, which might drive cysteine conversion to other metabolites. It was previously reported that global CDO1 deletion in mice caused multi-organ toxicity (7). Taurine supplementation did not prevent the pathology in global CDO1 KO mice, which was attributed to overproduction of H_2_S (7, 8). While liver may be less susceptible to CDO1 deletion compared to other organs in global CDO1 KO mice, liver mitochondrial impairment was also noted in the global CDO1 KO mice (7), suggesting that CDO1 knockout may also cause liver toxicity and organelle impairment that exacerbate liver injury upon additional metabolic stress. This may manifest as worsened liver injury and fibrosis under chronic fatty liver condition when CDO1 is only deleted in hepatocytes in mice. Cysteine metabolism is inter-connected with many other cellular pathways, and future studies are required to delineate the underlying drivers of liver injury in L-CDO1 KO mice.

Dietary cysteine has been linked to the development of obesity (14, 15, 19–21). In contrast, our study in L-CDO1-KO mice and L-CDO1-Tg mice shows that changes of liver CDO1-mediated cysteine catabolism, in either direction, does not have a significant impact on WD-induced obesity. These findings further imply that liver CDO1-mediated cysteine catabolism may have limited influence on circulating cysteine or system cysteine exposure. We also found that deletion of CDO1 in adipocytes did not affect WD-induced obesity, while CDO1 overexpression in adipocytes attenuated the development of obesity, which correlated with reduced hepatic steatosis. The mice were fed with WD that contained not only ∼42% fat calories but also 20% casein protein that provided adequate supply of dietary cysteine, which is potentially why a block of adipose cysteine catabolism did not further promote obesity development. In contrast, we have shown that reducing cysteine exposure in culture medium impaired adipogenesis in vitro (19), suggesting that reduced intracellular cysteine likely had a direct effect in inhibiting adiposity. In our mouse model, CDO1 is not only overexpressed in white adipocytes but also in BAT, which showed less fat accumulation in Ad-CDO1-Tg mice. Whether this is a result of reduced WAT or an intrinsic effect in brown adipocytes remains to be determined. Recent studies suggest that dietary cysteine restriction is associated with many metabolic changes including induction of FGF21, GSH and coenzyme A depletion, and sympathetic output, suggesting complex metabolic and neuronal changes in response to reduced cysteine intake. Interestingly, our previous study found that reduced dietary protein intake only reduced hepatic cysteine abundance without causing broad hepatic amino acid deficiency, a phenomenon that is not well understood (3). Short-term dietary protein restriction was associated with pronounced metabolic changes while long-term dietary protein restriction causes marked weight reduction (3, 19). These results imply that there might be a cysteine sensing mechanism that critically regulates the body’s metabolic responses, which remains to be further delineated in future studies.

Rare missense mutations of the CDO1 gene have been reported in humans who developed neurological disorders and developmental defects (24), suggesting that neuronal cells are sensitive to cysteine-mediated cytotoxicity. CDO1 gene has been found to be frequently silenced in various type of cancer cells and has been thought to play a tumor suppressor role (11, 25). By using genetic mouse models, we have shown that cysteine catabolism differentially modulates the susceptibility to diet-induced obesity and fatty liver disease, which warrants additional investigation to gain better understanding of cellular cysteine sensing mechanisms and cysteine control of cell metabolism. Whether genetic variations and epigenetic regulation of CDO1 may modulate metabolic disease risk is currently unclear and remains to be investigated in the future.

## Methods

### Reagents

Anti-CDO1 antibody (12589-1-AP) was purchased from Proteintech Group, Inc. (Rosemont, IL). Anti-Actin antibody (A5441) was purchased from Sigma (St. Louis, MO). α-Tubulin antibody (ab6046) were purchased from Abcam (Cambridge, MA). Aspartate aminotransferase (AST) kit, alanine aminotransferase (ALT) assay kit, total triglyceride assay kit, and total cholesterol assay kit were purchased from Pointe Scientific (Canton. MI). AAV8-TBG-Null and AAV8-TBG-cre were purchased from Vector Biolabs (Malvern, PA).

### Mice

As described previously (10), the *Cdo1* flox mice on C57BL/6N genetic background were developed with CRISPR/Cas9 technology by Cyagen Biosciences Inc. (Santa Clara, CA). Cre recombinase-mediated deletion of a 2074 bp region contains the entire exon 3 of the mouse *Cdo1* gene. AAV-TBG-cre (2 x 10^11^ GC/mouse) was injected to *Cdo1* flox mice via tail vein to delete the *Cdo1* gene in hepatocytes. WT controls were *Cdo1* flox mice injected with AAV8-TBG-Null. *Cdo1* flox mice were crossed with Adipoq-Cre mice (strain #: 028020) on C57BL/6J background purchased from the Jackson Laboratory (Bar Harbor, ME). WT controls were homozygous *Cdo1* flox littermates without Adipoq-cre. The *Cdo1*(*Rosa26*) knock-in (KI) mice on C57BL/6NTac genetic background were generated by inserting a “CAG promoter-loxP-3*SV40 pA-loxP-Kozak-mouse Cdo1 CDS-rBG pA” cassette containing the mouse *Cdo1* cDNA sequence into the Intron 1 of *Rosa26* in reverse orientation using CRISPR technology (Cyagen Biosciences Inc.). Homozygous *Cdo1*(*Rosa26*) KI mice were injected 2×10^11^ GC/mouse AAV8-TBG-cre (L-CDO1-Tg) or AAV8-TBG-Null (WT controls) via tail vein. Ad-CDO1-Tg mice were generated by crossing *Cdo1*(*Rosa26*) KI mice with Adipoq-cre mice. Ad-CDO1-Tg mice were homozygous for *Cdo1(Rosa26)* and heterozygous for Adipoq-cre. WT controls were homozygous *Cdo1*(*Rosa26*) KI littermates without Adipoq-cre. Mice were housed in micro-isolator cages with Biofresh performance bedding (Pelleted cellulose) under 7 am - 7 pm light cycle and 7 pm −7 am dark cycle. Western diet contains 42% fat calories and 0.2% cholesterol was previously described (19). The HFCFr diet was previously described (23). Animals received humane care according to the criteria outlined in the “Guide for the Care and Use of Laboratory Animals.” Mice were euthanized with isoflurane inhalation before tissue collection. All animal protocols were approved by the Institutional Animal Care and Use Committee of the University of Oklahoma Health (Protocol # 22–072).

### Liver lipid measurement

Liver lipids were extracted from liver tissues with a mixture of chloroform:methanol (2:1; volume:volume), dried under nitrogen, and resuspended in isopropanol containing 1% triton X-100. Liver triglyceride and cholesterol were measured by using colorimetric assay kits.

### Western Blotting

Tissues were homogenized in radioimmunoprecipitation assay (RIPA) buffer (150 mM NaCl, 50 mM Tris-HCl, pH 7.4; 1% NP-40; 0.25% sodium deoxycholate; 1 mM EDTA; protease inhibitor cocktail), incubated on ice for 1 hour, and centrifuged at 13,000 *× g* for 15 min at 4 °C. Clear lysates were mixed with equal volume of 2X Laemmli buffer and heated at 95 °C for 5 minutes and used for SDS-PAGE and immunoblotting.

### Real-time PCR

Total RNA was isolated using TRIzol Reagent (15596026, ThermoFisher Scientific, Waltham, MA). Total RNA was used for cDNA synthesis using Oligo dT primer and SuperScript III reverse transcriptase (18080093, ThermoFisher Scientific, Waltham, MA). Real-time PCR was performed with iQ SYBR Green Supermix (1708882, Bio-Rad Laboratories, Inc., Hercules, CA) on a BioRad CFX384 Real-Time System. Relative mRNA expression is calculated using the comparative Ct (Ct) method and expressed as 2^−ΔΔCt^. 18s or GAPDH was used as internal control for normalization.

### Hematoxylin-eosin, Sirius red, and immunohistochemistry (IHC) staining

Adipose and liver tissues were fixed in 4% paraformaldehyde at 4 °C for 24 h. Tissues were then embedded and cut for hematoxylin-eosin stain and Sirius red stain. For IHC stain of CDO1 and F4/80, tissue sections were deparaffinized and hydrated. Antigen retrieval was done by boiling slides in citric acid buffer (10 mM citric acid, 0.05% Tween-20, pH 6.0) for 20 minutes. Slides were then blocked with 5% normal goat serum + 3% BSA followed by overnight incubation with primary antibody. Endogenous peroxidase activity was blocked by 0.3% hydrogen peroxide. The slides were then incubated with secondary antibody and HRP substrate. Images were acquired with a ThermoFisher EVOS M5000 Imaging System (Waltham, MA).

### Liver metabolite measurement

Liver metabolites were measured as part of the unbiased metabolomics performed by Metabolon Inc (Durham, NC) as previously described (3). Metabolon’s reference library and software were used to process identified peaks from raw data. The analysis does not include standard curve. Area-under-the-curve (AUC) of peaks were calculated. Controls were arbitrarily set as “1”.

### Statistical analysis

**All results are expressed as** mean ± SEM. The p values were calculated using either unpaired t-test or 2-way ANOVA and post hoc tests. A p value < 0.05 is considered significant. “*”, < 0.05; “**”, < 0.01; “***”, < 0.001; “****”, < 0.0001.

## Acknowledgement

supported by NIH grants 1R01 DK117965-01A1 and 1R01DK131064-01

## References

1. M. H. Stipanuk, J. E. Dominy, Jr., J. I. Lee, R. M. Coloso, Mammalian cysteine metabolism: new insights into regulation of cysteine metabolism. J Nutr 136, 1652S–1659S (2006).

2. T. Li, J. Y. L. Chiang, Bile Acid Signaling in Metabolic and Inflammatory Diseases and Drug Development. Pharmacol Rev 76, 1221–1253 (2024).

3. D. Matye et al., TFEB regulates sulfur amino acid and coenzyme A metabolism to support hepatic metabolic adaptation and redox homeostasis. Nat Commun 13, 5696 (2022).

4. L. L. Hirschberger, S. Daval, P. J. Stover, M. H. Stipanuk, Murine cysteine dioxygenase gene: structural organization, tissue-specific expression and promoter identification. Gene 277, 153–161 (2001).

5. H. Jurkowska et al., Primary hepatocytes from mice lacking cysteine dioxygenase show increased cysteine concentrations and higher rates of metabolism of cysteine to hydrogen sulfide and thiosulfate. Amino Acids 46, 1353–1365 (2014).

6. J. E. Dominy, Jr., L. L. Hirschberger, R. M. Coloso, M. H. Stipanuk, Regulation of cysteine dioxygenase degradation is mediated by intracellular cysteine levels and the ubiquitin-26 S proteasome system in the living rat. Biochem J 394, 267–273 (2006).

7. I. Ueki et al., Knockout of the murine cysteine dioxygenase gene results in severe impairment in ability to synthesize taurine and an increased catabolism of cysteine to hydrogen sulfide. Am J Physiol Endocrinol Metab 301, E668–684 (2011).

8. H. B. Roman et al., The cysteine dioxgenase knockout mouse: altered cysteine metabolism in nonhepatic tissues leads to excess H2S/HS(-) production and evidence of pancreatic and lung toxicity. Antioxid Redox Signal 19, 1321–1336 (2013).

9. Y. Wang, et al., Bile acids regulate cysteine catabolism and glutathione regeneration to modulate hepatic sensitivity to oxidative injury. JCI Insight 3 (2018).

10. J. Chen et al., Deletion of hepatocyte cysteine dioxygenase type 1, a bile acid repressed gene, enhances glutathione synthesis and ameliorates acetaminophen hepatotoxicity. Biochem Pharmacol 222, 116103 (2024).

11. Y. P. Kang et al., Cysteine dioxygenase 1 is a metabolic liability for non-small cell lung cancer. Elife 8 (2019).

12. R. Leonardi, J. E. Rehg, C. O. Rock, S. Jackowski, Pantothenate kinase 1 is required to support the metabolic transition from the fed to the fasted state. PLoS One 5, e11107 (2010).

13. A. K. Elshorbagy, V. Kozich, A. D. Smith, H. Refsum, Cysteine and obesity: consistency of the evidence across epidemiologic, animal and cellular studies. Curr Opin Clin Nutr Metab Care 15, 49–57 (2012).

14. L. El-Khairy, P. M. Ueland, O. Nygard, H. Refsum, S. E. Vollset, Lifestyle and cardiovascular disease risk factors as determinants of total cysteine in plasma: the Hordaland Homocysteine Study. Am J Clin Nutr 70, 1016–1024 (1999).

15. L. El-Khairy, S. E. Vollset, H. Refsum, P. M. Ueland, Predictors of change in plasma total cysteine: longitudinal findings from the Hordaland homocysteine study. Clin Chem 49, 113–120 (2003).

16. T. Olsen et al., Dietary sulfur amino acid restriction in humans with overweight and obesity: a translational randomized controlled trial. J Transl Med 22, 40 (2024).

17. S. N. Nichenametla et al., Cysteine restriction-specific effects of sulfur amino acid restriction on lipid metabolism. Aging Cell 21, e13739 (2022).

18. A. K. Elshorbagy et al., Cysteine supplementation reverses methionine restriction effects on rat adiposity: significance of stearoyl-coenzyme A desaturase. J Lipid Res 52, 104–112 (2011).

19. D. J. Matye, H. Wang, Y. Wang, L. Xiong, T. Li, Bile acid sequestrant inhibits gluconeogenesis via inducing hepatic cysteine dioxygenase type 1 to reduce cysteine availability. Am J Physiol Gastrointest Liver Physiol 328, G166–G178 (2025).

20. A. H. Lee et al., Cysteine depletion triggers adipose tissue thermogenesis and weight loss. Nat Metab 7, 1204–1222 (2025).

21. A. Varghese et al., Unravelling cysteine-deficiency-associated rapid weight loss. Nature 643, 776–784 (2025).

22. J. Niewiadomski et al., Effects of a block in cysteine catabolism on energy balance and fat metabolism in mice. Ann N Y Acad Sci 1363, 99–115 (2016).

23. D. J. Matye et al., Combined ASBT Inhibitor and FGF15 Treatment Improves Therapeutic Efficacy in Experimental Nonalcoholic Steatohepatitis. Cell Mol Gastroenterol Hepatol 12, 1001–1019 (2021).

24. L. Rowe et al., A proposed role for CDO1 in CNS development: Three children with rare missense variants and a neurological phenotype. HGG Adv 6, 100417 (2025).

25. J. Jeschke et al., Frequent inactivation of cysteine dioxygenase type 1 contributes to survival of breast cancer cells and resistance to anthracyclines. Clin Cancer Res 19, 3201–3211 (2013).

